# How reliable are ligand-centric methods for Target Fishing?

**DOI:** 10.1101/032946

**Authors:** Antonio Peón, Cuong C. Dang, Pedro J. Ballester

## Abstract

Computational methods for Target Fishing (TF), also known as Target Prediction or Polypharmacology Prediction, can be used to discover new targets in small-molecule drugs. This may result in repositioning the drug in a new indication or improving our current understanding of its efficacy and side effects. While there is a substantial body of research on TF methods, there is still a need to improve their validation, which is often limited to a small part of the available targets and not easily interpretable by the user. Here we discuss how target-centric TF methods are inherently limited by the number of targets that can possibly predict (this number is by construction much larger in ligand-centric techniques). We also propose a new benchmark to validate TF methods, which is particularly suited to analyse how predictive performance varies with the query molecule. On average over approved drugs, we estimate that only five predicted targets will have to be tested to find two true targets with submicromolar potency (a strong variability in performance is however observed). In addition, we find that an approved drug has currently an average of eight known targets, which reinforces the notion that polypharmacology is a common and strong event. Furthermore, with the assistance of a control group of randomly-selected molecules, we show that the targets of approved drugs are generally harder to predict.

## 1. Introduction

Target Fishing (TF)(Cereto-Massagué et al., 2015; Lavecchia and Cerchia, 2015), also known as Target Prediction or Polypharmacology Prediction, consists in predicting the macromolecular targets of a query molecule. This problem is the reverse of Virtual Screening (VS) (Schneider, 2010; Sukumar and Das, 2011), where the goal is to predict the ligands of a query target. Computational methods for TF are of great interest, as identifying previously unknown targets of a molecule is the basis of a number of important drug design and chemical biology applications (Ursu and Waldmann, 2015). Indeed, discovering a new target in a drug could lead to its reposition in a new indication as well as an enhanced understanding of its efficacy and side-effects (Huang et al., 2014). Furthermore, these tools can be used for target deconvolution of phenotypic screening hits (Lee and Bogyo, 2013), which is a prerequisite to gain mechanistic understanding of phenotypic activity and helpful for drug development. This two-stage process, phenotypic screening followed by target deconvolution, constitutes an attractive alternative strategy for the discovery of molecularly targeted therapies.

The fast growth of freely-available bioactivity resources, e.g. PubChem (Cheng et al., 2014) or ChEMBL (Bento et al., 2014), has sparked a new generation of powerful data-driven methods for TF. This growth is exemplified by the ChEMBL database, which in a few years has assembled and fully curated chemical structures and bioactivities from more than 50,000 scientific publications (Bento et al., 2014). Moreover, this database is periodically updated and will eventually incorporate a flood of new data that is being extracted from the patent literature (Papadatos et al., 2015). As discussed by (Cereto-Massagué et al., 2015), TF methods have been categorised into those based on molecular similarity (Liu et al., 2014), machine learning (van Laarhoven et al., 2011), protein structure analysis (Gao et al., 2008) and bioactivity spectra analysis (Füllbeck et al., 2009; Holbeck et al., 2010). Some of these methods have been made available as web servers (Gfeller et al., 2014; Wang et al., 2013).

Here we propose a new classification of TF methods into two broad categories: *target-centric* and *ligand-centric*. Target-centric methods are defined as those building a predictive model for each considered target. Each of these models is thereafter used to predict whether the query molecule has activity against the corresponding target (other names for the query molecule are common, such as test compound, test molecule or test ligand). Thus, this panel of models provides a set of predicted targets for any query molecule. Many target-centric TF methods are based on multi-target Quantitative Structure–Activity Relationship (QSAR) models (Speck-Planche and Cordeiro, 2015; Zanni et al., 2014). The model is typically trained on large sets of active and inactive target-ligand instances derived from a database of target-annotated molecules (the training set). These models have employed various regression or classification techniques, such as Kernel Classifiers (van Laarhoven et al., 2011), Winnow (Nigsch et al., 2008), Ranking Perceptron (Yu et al., 2012), Random Forest (Yu et al., 2012) or Naïve Bayes Classifier (Koutsoukas et al., 2013). Instead of supervised learning, other target-centric techniques are based on unsupervised learning such as the Similarity Ensemble Approach (SEA)(Keiser et al., 2007, 2009). SEA constructs a model for each target estimating how likely is the query molecule to belong to the set of cognate ligands of the target based on an underlying molecular similarity metric. In addition, there are target-centric methods that can estimate whether the query molecule binds to a structural model of the target (Schomburg and Rarey, 2014). These methods are in principle able to interrogate targets without known ligands, although their success largely depends on the accuracy of the employed scoring function (Ain et al., 2015).

On the other hand, ligand-centric methods are those based on the similarity of the query molecule to a very large set of target-annotated molecules. This similarity can be in terms of 2D chemical structure (Nettles et al., 2006) using circular fingerprints (Rogers and Hahn, 2010), 3D molecular properties (Cortés-Cabrera et al., 2013) using Ultrafast Shape Recognition variants (Armstrong et al., 2010; Ballester and Richards, 2007; Ballester, 2011) or NCI-60 bioactivity spectra (Holbeck et al., 2010) using cellular fingerprints (Füllbeck et al., 2009). Note that there are similarity-based methods that are not ligand-centric. This is the case of TAMOSIC (Wang et al., 2013), which learns the optimal similarity cutoff for each target with at least 30 ligands.

An important advantage of ligand-centric methods over target-centric methods has been so far overlooked. Whereas ligand-centric methods can interrogate any target that has at least one known ligand, target-centric models can only evaluate the typically much smaller set of targets for which a model can be built. For instance, TarFisDock (Gao et al., 2008) predictions are limited to 1100 targets with available crystal structure and known binding site, whereas SEA (Keiser et al., 2007, 2009) only evaluates targets with at least five known ligands. This means that target-centric methods are by construction blind to up to thousands of targets considered by ligand-centric techniques, but this is not obvious as target-centric performance is only evaluated on qualifying targets. Target-centric and ligand-centric methods are nevertheless complementary. Indeed, in cases where the targets of interest are known to have many ligands, more accurate target-centric models could be possible and thus these tools are likely to be more suitable. By contrast, in cases where evaluating as many targets as possible is preferable, ligand-centric tools would be more appealing, as these provide a much wider coverage of the proteome.

Unfortunately, it is unclear how well ligand-centric methods work in practice due to the limitations of existing benchmarks. Some validations have been restricted to a few tens of ligand-rich targets using benchmarks borrowed from VS (AbdulHameed et al., 2012) and thus tell us very little about how well the methods will perform on the many remaining targets. Furthermore, some performance measures, such as the ROC (Receiver Operating Characteristic) AUC (Area Under Curve), do not precisely measure TF performance. For example, how many true targets of a query molecule one is likely to find in practice using a method that has obtained an average ROC AUC of 0.7 over 40 targets? On the other hand, TF is often posed as a multi-category classification problem, which formulates a binary classification problem per target and thus the variation of predictive performance across query molecules has not been analysed in these studies. Importantly, these benchmarks exclude many possible targets of the analysed molecules because the corresponding target-centric models could not be trained on the excluded targets. As a result of these limitations, current benchmarks offer little guidance on pragmatic questions such as how many predicted targets have to be tested on average to find a true target, how many known targets are typically missed or how such performance varies with the query molecule.

In this study, we propose a new benchmark to validate TF methods, which naturally lends itself to answer such questions. This is based on formulating a binary classification problem for each query molecule. From this new perspective, we provide a lower-bound for the current performance of ligand-centric methods representing the minimum that can be expected nowadays from them. As a byproduct, our analysis provides an update for the degree of polypharmacology observed in approved drugs. The rest of the paper is organised as follows. Section 2 describes the experimental setup, including data selection, data partitions, TF method and performance metrics. Section 3 discusses the results. Section 4 presents the conclusions.

## 2. Experimental setup

This section describes the setup of all the numerical experiments carried out in this study. This setup is composed of the following elements: data selection, data partitions, TF method and measures of predictive performance. All molecular data processing is done with SQL queries from Python 2.7.9 on a local copy of the ChEMBL database running PostgreSQL 9.4.3, with molecular similarity searches using in addition the RDKit PostgreSQL cartridge (2015.03.1 release).

### 2.1. Data selection

The first step is constructing datasets from the ChEMBL database. We started by downloading release 20 as a PostgreSQL dump (ChEMBL_20 release), which contains data for 10,774 targets, 1,456,020 molecules with disclosed chemical structure and 13,520,737 bioactivities.

Single-protein was the most common target type (6,018 of the 10,774 targets). In order to provide the most specific target prediction, we restricted to single-protein targets, which incidentally constitutes the largest molecular target type in the database (the „protein complex’, „protein family’ and „nucleic-acid’ types only have 261, 217 and 29 targets, respectively). The remaining general constraints were requiring the maximum confidence_score = 9 (i.e. direct single-protein target assigned by the data curator), activities.published_relation = '=', assay_type = 'B' and standard_units = 'nM'. As a result of this process, 888,354 molecules were found to be associated to the 6,018 single-protein targets through 4,871,527 bioactivities.

Further requirements are commonly imposed for the measured bioactivity of a ligand against a target to be counted as a known target for that ligand. First, the bioactivity measurement must be of relatively high quality, activities.standard_type IN ('EC50','Ki','Kd','IC50'), which discards percentages of inhibition among other lower-quality measurements. Second, only complexes with a sufficiently potent bioactivity are retained (common activity thresholds are 1μM and 10μM meaning that a ligand hitting any target with an activity higher than 10μM will not be considered to be a target in neither of these two scenarios). Third, only targets with at least n qualifying ligands are considered. For many target-centric methods, a sufficiently high number of ligands is needed to build a model for the target, e.g. those methods based on similarity-ensemble approaches (n=5) (Keiser et al., 2009) or multi-target QSAR (n=20) (Koutsoukas et al., 2013). In this study, we analyse ligand-centric methods, which can evaluate any target with at least a known ligand (i.e. n=1) and hence result in a much broader search for targets (3,035 molecular targets with 10μM). An analysis of the target coverage of TF methods is carried out in section 3.1.

### 2.2. Data partitions

Next, we partition each of the two n=1 datasets as follows. First, we identify the subset of approved drugs. Second, we search for all those approved drugs in the ChEMBL database meeting the criteria, with a suitable chemical structure available and hitting any of the targets introduced in the previous section. These are the two approved-drugs sets of query molecules shown in Table 1. Third, we pick at random two further sets of molecules of the same size, which we called random-molecules sets. This will serve as a control group to investigate how target predictions for marketed drugs differ from those made for other types of molecules. The rest of ligands forms the set of database molecules, which is the same for both sets of query molecules but different between thresholds. Table 1 shows the four non-overlapping data partitions A-D (no query molecule is included as database molecule too).

**Table 1.** Dataset size depending on modelling constraints.

| ID | Dataset | #query-molecules | #database-molecules |
| --- | --- | --- | --- |
| A | approved-drugs_Thres10μM | 745 | 183,282 |
| B | random-molecules_Thres10μM | 745 | 183,282 |
| C | approved-drugs_Thres1μM | 617 | 147,027 |
| D | random-molecules_Thres1μM | 617 | 147,027 |

### 2.3. A simple TF method to estimate a lower-bound for performance

For our analysis, we selected a simple two-dimensional chemical similarity search (Willett, 2014) in order to obtain a lower-bound for the performance of ligand-centric TF methods. This goal requires selecting a simple method, rather than an optimal method which would be unlikely to provide such lower-bound. Consequently, we selected the dice score on MACCS fingerprints *ad hoc,* although there are of course other valid choices too. We started by generating MACCS fingerprints (Durant et al.) for all query and database molecules in Table 1. Each fingerprint encodes the presence or absence of 166 predetermined chemical groups in the molecule as a binary string of the same size. These were generated using the RDKit (Lamdrum).

As usual, fingerprints could not be generated for a few unusual molecules and consequently queries could not be performed for these. This is the case of Gramidicin (CHEMBL1201469), which is actually not a molecule but a mixture of three antibiotic compounds. Other examples are some organometallic compounds such as the anti-rheumatic agent Auranofin (CHEMBL1366). Table 1 compiles all selected molecules for which MACCS fingerprints could be generated.

Using their MACCS fingerprints, the Dice score was used to measure the similarity between a query molecule and all the database molecules. The Dice score is defined as:

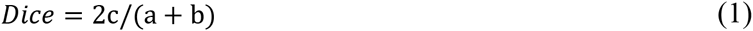

where a is the number of on bits in molecule A, b is number of on bits in molecule B, while c is the number of bits that are on at the same positions in both molecules. For each query, the top k hits can be identified from the corresponding ranking of database molecules (these are the k database molecules with the most similar chemical structure to that of the query molecule). We consider here k = 1, 5, 10 and 15 to investigate the dependence of the method with its only control parameter k.

Finally, the known targets for the k hits are retrieved from the ChEMBL database and returned as predicted targets for the considered query molecule. Thus, a set of predicted targets is obtained for each combination of query molecule and k value. Note that a known target is not just any target annotated in the ChEMBL database, but one complying with the requirements set in section 2.1. for each of the four cases in Table 1.

### 2.4. Measuring predictive performance

Each performed query can be posed as a separate classification problem. For validation purposes, the known targets of the query molecule are taken as a ground truth. Thus, we assume that the known targets are all the qualifying targets of the molecule, whereas the rest of considered targets are non-targets for that molecule. However, as the query molecule has only been tested against less than 0.1% of the ChEMBL targets on average, it is expected that many unconfirmed targets, especially those coming from molecules similar to the query molecule, would be actually targets if only these could be comprehensively tested. As a result, any empirically untested target-ligand association that is predicted to be a true association will have to be rejected as false, despite an unknown part of these being actually true targets of the molecule. We must therefore keep in mind that this retrospective validation represents a lower-bound for performance in this sense as well.

Table 2 shows the confusion matrix arising from assessing target predictions against experimental evidence for each query molecule. After the assessment, each target prediction can be classed in one of four categories: TP for True Positive (the predicted target is a known target); TN for True Negative (the target was not predicted but anyway is not known to be a target); FP for False Positive (the predicted target is not known to be a target, i.e. a false discovery or Type I error); and FN for False Negative (the target was not predicted and it is actually a target, i.e. missed discovery or Type II error).

**Table 2.** Confusion matrix arising from assessing target predictions against experimental evidence for each query molecule.

| Target | Predicted | Non-predicted |
| --- | --- | --- |
| **Yes** (experimentally tested) | TP | FN |
| **No** (not tested/tested) | FP | TN |

From these quantities, we will calculate four performance measures per query molecule. Accuracy is the proportion of correct target predictions:

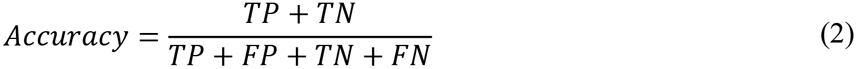

Precision is the proportion of new targets that would be obtained after experimentally validating the predictions of the method:

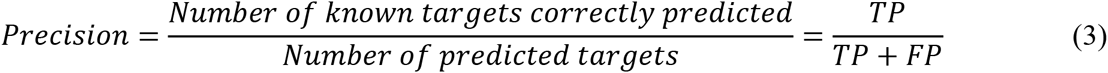

Recall accounts for the proportion of true targets that the method has missed:

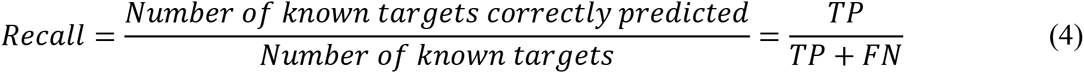

The Matthews Correlation Coefficient (MCC) captures both types of error in a single metric, with higher values being better up to +1 (perfect classification):

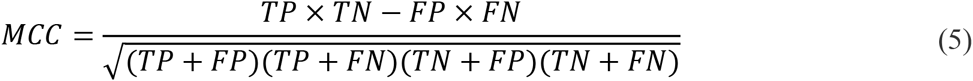

Lastly, the Number of Predicted Targets (NPT) will be also reported to investigate how this varies with the method’s control parameter k. The entire workflow is sketched in Figure 1.

**Figure 1.**
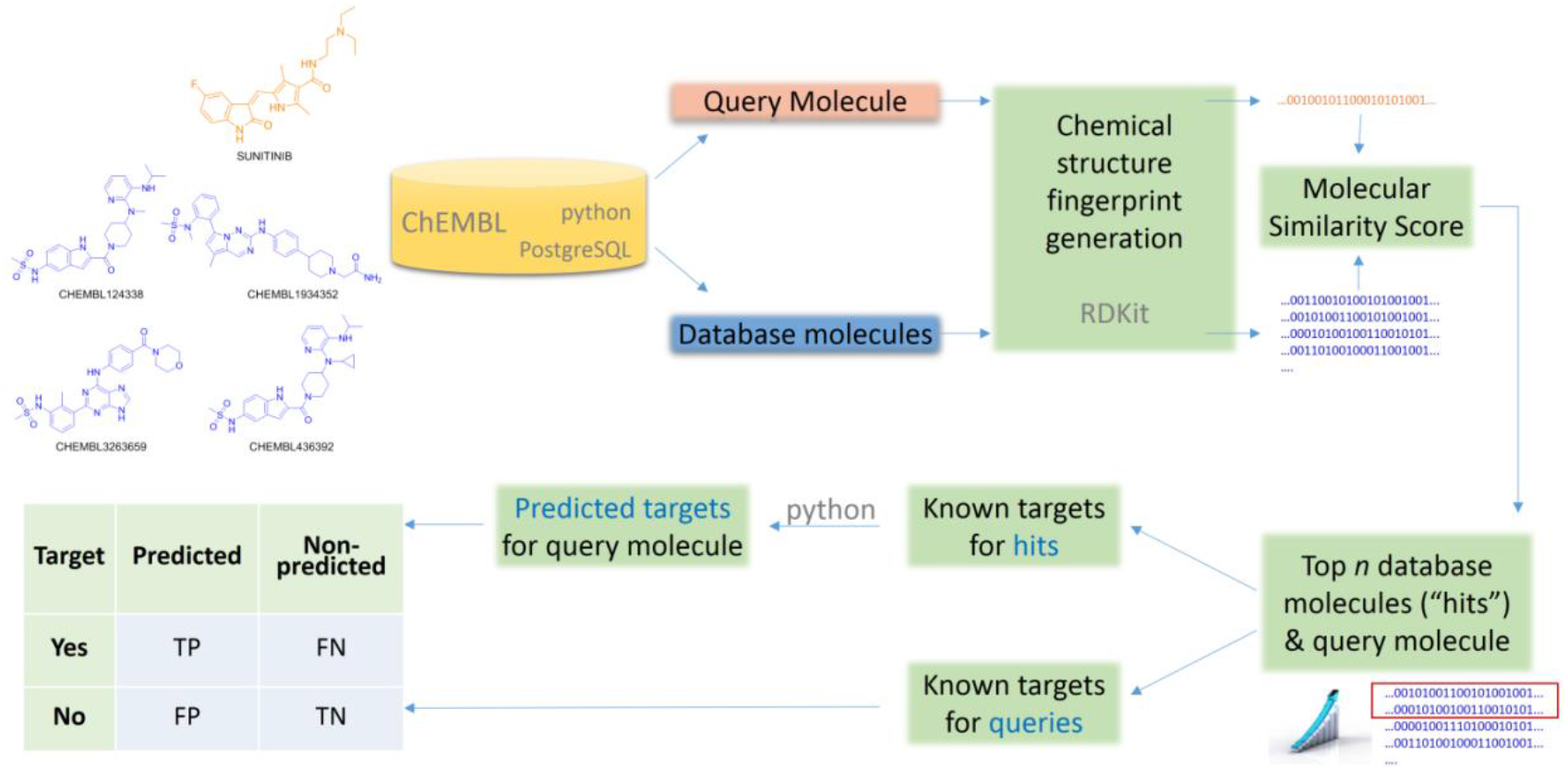
Generic workflow to apply and validate a ligand-centric method for TF.

## 3. Results and Discussion

Four key questions along with two representative case studies are addressed in this section. The analysis is based on the performance obtained by the query molecules in the four datasets in Table 2, which will be summarised with boxplots of precision, recall, MCC and NPT.

### 3.1. How many targets are being missed by target-centric TF techniques?

The first two rows of Table 3 show the number of targets considered by a ligand-centric TF method with two target definitions (i.e. activity thresholds of 1μM and 10μM). The remaining rows show the number of targets considered by exemplary target-centric methods as a result of only considering targets with at least 5-40 ligands. To allow a fair comparison, we have calculated the number of targets using the same selection criteria on chembl20 data (section 2.1), except for the minimum number of ligands required by each method and the selected activity threshold.

**Table 3.** Numbersof considered targets and number of missed targets depending on the data selection criteria of the employed benchmark.

| Dataset | Only targets with at least | & activity below | #Targets in study | #Targets if chembl20 | #Targets missed |
| --- | --- | --- | --- | --- | --- |
| This study | 1 ligand | 1μM | 2,580 | 2,580 | 0 |
| This study | 1 ligand | 10μM | 3,035 | 3,035 | 0 |
| (Keiser et al., 2007) | 5 ligands | 1μM | 246 | 1,788 | 792 |
| (Mugumbate et al., 2015) | 10 ligands | 10μM | 1,543 | 1,804 | 1,231 |
| (Koutsoukas et al., 2013) | 20 ligands | 10μM | 894 | 1,378 | 1,657 |
| (Wang et al., 2013) | 30 ligands | 10μM | 794 | 1,104 | 1,931 |
| (Martínez-Jiménez et al., 2013) | 40 ligands | 10μM | 1,258 | 917 | 2,118 |

For example, target-centric methods powered by models requiring at least 40 ligands per target and defining a target with an activity threshold of 10μM would be predicting whether the query molecule has activity against any of the 917 qualifying single-protein targets. In contrast, a ligand-centric method with the same activity threshold will be able to evaluate 2,118 targets more, for which the first method is unable to provide any prediction by construction. Of course, the advantage of target-centric over ligand-centric methods is that the former will tend to perform better on those targets with a high number of ligands, which highlights the complementarity of both approaches. It would be interesting if the performance of target-centric methods was evaluated per target and analysed against its number of cognate ligands, as it is currently unknown how reliable are their predictions on the many targets with only a few ligands above the minimum.

### 3.2. How many targets are typically hit by a molecule?

For each of the four cases in Table 2, Figure 2 shows boxplots summarising the distribution of the number of known single-protein targets (NKTs) across query molecules.

**Figure 2.**
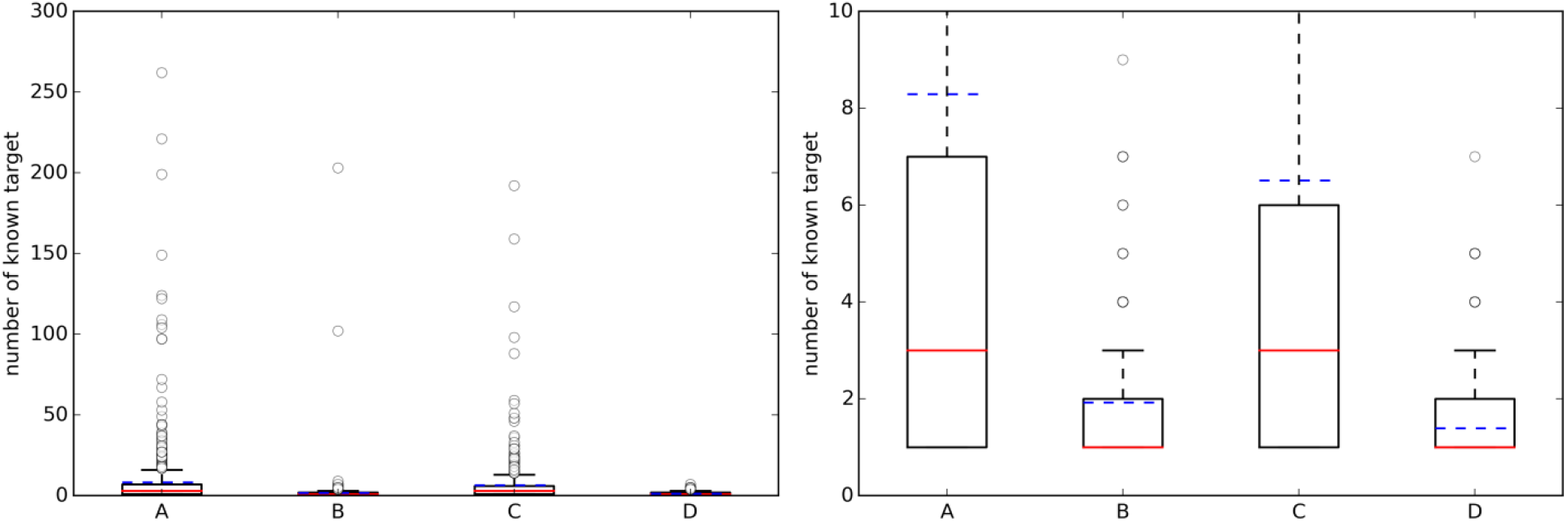
(**left**) Boxplots with the number of known targets (NKT) across query molecules. (**right**) A zoom of the same boxplots on the right (the average number of known targets is marked with a dashed blue line, whereas the median is given by the continuous red line). A = (Approved, 10μM), B = (Random, 10μM), C = (Approved, 1μM), D = (Random, 1μM).

On the left, a substantial number of strong outliers are appreciated. These correspond to promiscuous query molecules such as sunitinib, which has 192 submicromolar targets (262 targets using the 10 μM threshold). In contrast, there are also seemingly selective drugs like the antiretroviral agent Nelfinavir with only one known target below 1μM (HIV-1 protease; although there are also many non-molecular targets annotated in ChEMBL for this drug). On the right, we can appreciate that approved drugs currently have an average of eight known targets with potency better than 10μM, although the median number is three targets. This new estimate is based on 745 drugs and their 1,076 targets and it is two targets higher than previous estimates using less data (802 drugs and 480 targets) (Mestres et al., 2009). However, the boxplot’s lower quartile value indicates that at least 25% of these drugs have just one known target and thus seem very selective. It is also noteworthy in Figure 2 that the number of annotated targets for the set of random molecules is smaller than that for approved drugs, with four targets on average instead of eight. This substantial difference is likely to be due to a much higher number of targets being tested during the process of developing a drug.

### 3.3. How many predicted targets have to be tested to find a true target?

Table 4 presents average performance results for approved drugs (set A), with the TF method using four different k values. As k increases, Type I errors increase (lower precision) and Type II errors decrease (higher recall). In other words, as more top hits are used to provide predicted targets, fewer known targets are missed. However, this comes at the cost of having more false positives, as target inferences are made using increasingly less similar database molecules. Using the top 5 hits to predict targets (i.e. k=5) provides the best compromise between these conflictive objectives (i.e. the highest average MCC). This setting leads to 5.04 predicted targets on average over these query molecules (note that each top hit may have more than one known target, but collectively provide fewer targets because some of these are repeated in the set). Lastly, the very high average accuracy values are due to each classification problem being highly unbalanced and the method correctly discarding the vast majority of non-targets. Nevertheless, unlike precision and recall, accuracy is not suitable to measure Type I and II errors and hence is not helpful to address the investigated questions.

**Table 4.** Average (av) performance of the TF method on query molecules from set A= (Approved, 10μM). These molecules have an average of 8.3 known targets.

| k | avNPT | avAccuracy | avPrecision | avRecall | avMCC |
| --- | --- | --- | --- | --- | --- |
| 1 | 1.92 | 0.997 | 0.434 | 0.212 | 0.269 |
| 5 | 5.04 | 0.997 | 0.352 | 0.341 | 0.303 |
| 10 | 7.91 | 0.996 | 0.296 | 0.403 | 0.300 |
| 15 | 10.32 | 0.995 | 0.257 | 0.437 | 0.289 |

Figure 3 shows the distribution of the NPT across query molecules using k=5. By comparing it with Figure 2, it is observed that there are substantially more known targets than predicted targets for approved drugs using the top 5 hits for predictions (this is not the case for the sets of random molecules, where most query molecules have a higher number of predicted targets than of known targets).

**Figure 3.**
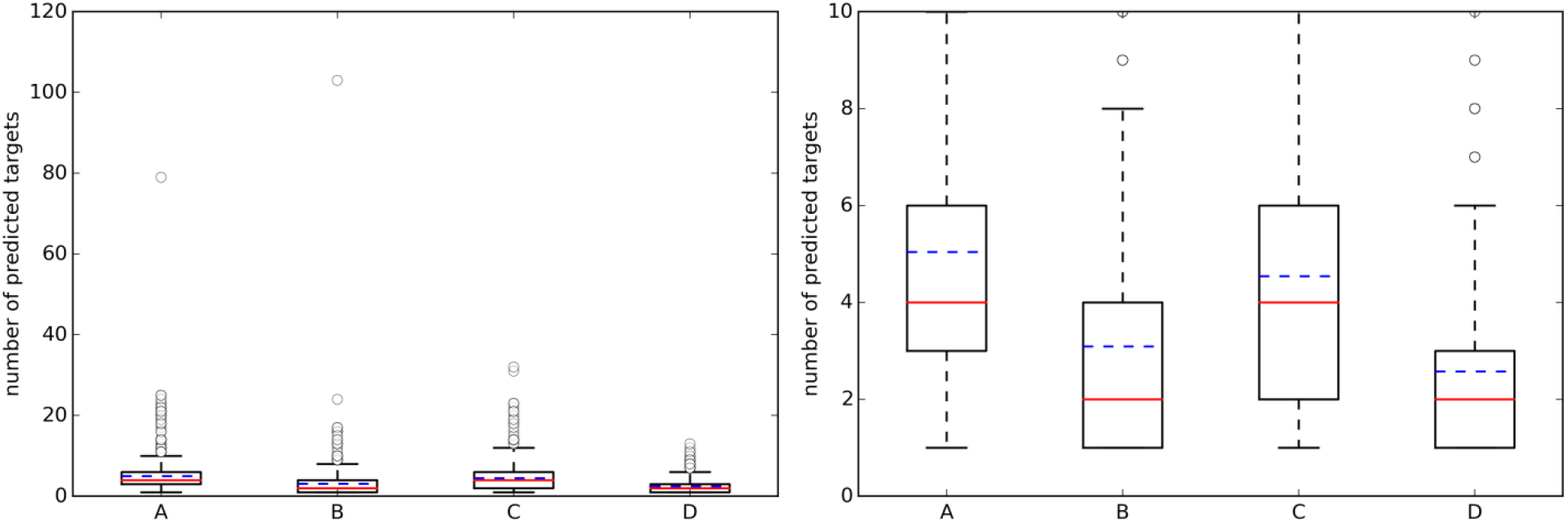
(**left**) Boxplots with the number of predicted targets (NPT) across query molecules using k=5; (**right**) A zoom of the same boxplots on the right (the average number of known targets is marked with a dashed blue line, whereas the median is given by the continuous red line). A = (Approved, 10μM), B = (Random, 10μM), C = (Approved, 1μM), D = (Random, 1μM).

Figure 4(left) summarises the distribution of precision results across the query molecules. For approved drugs, the average precision is 0.35 in the 10μM case (0.38 in the 1μM case). That is, despite the simplicity of the method and thanks to the wealth of data on which it relies, only five predicted targets will have to be tested in order to find two true targets with potency better than 1μM. In all cases, there is strong performance variability across the query molecules, as it can be appreciated by the large interquartile range of each boxplot. For instance, in set C, the predictions for the targets of 109 drugs are of the highest precision (precision=1), those for other 216 drugs not precise at all (precision=0) and those for the remaining 420 drugs have intermediate precision values (in other words, hit rates are neither 0% nor 100%). Also, the cases with a tighter activity threshold of 1μM are on average slightly better predicted than their counterparts using 10μM. Specific cases with high- and low-precision performance will be discussed in section 3.5. On the other hand, the sets with random molecules obtained much better results than those with approved drugs. Thus, if we order the four cases by average precision (dashed blue line in Figure 4), this gives the following performance hierarchy D>B>C>A (i.e. D obtains higher average precision than B, B better than C and C better than A). Interestingly, this is the opposite ranking for the number of known targets (A>C>B>D). In other words, those sets with a higher number of known targets tend to be harder to predict.

**Figure 4.**
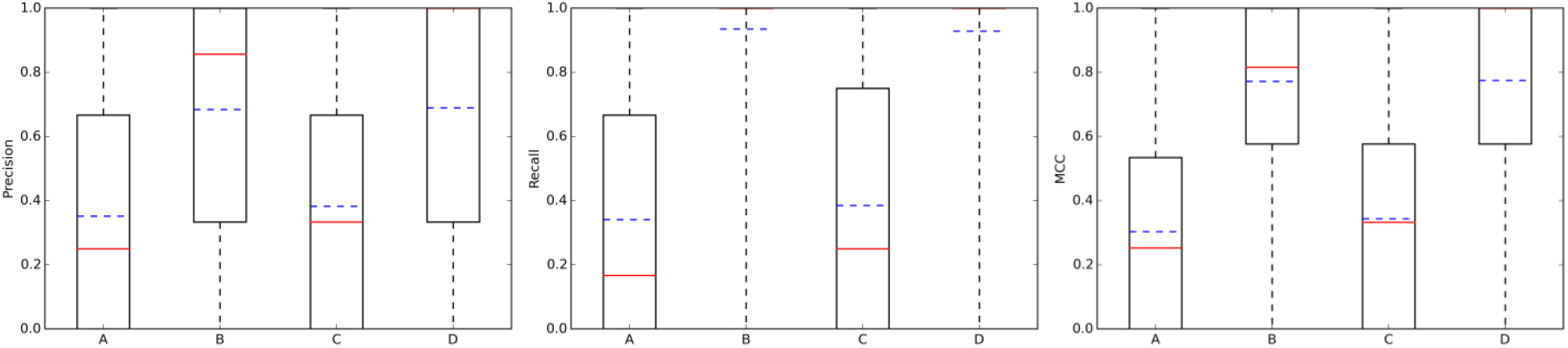
Performance the TF method with k=5. From left to right, boxplots for precision, recall and mcc across the query molecules in each of the four data partitions: A = (Approved, 10μM), B = (Random, 10μM), C = (Approved, 1μM), D = (Random, 1μM). The average and median values of each performance metric are shown as dashed blue lines and continuous red lines, respectively.

However, the cause of obtaining lower predictive accuracy with approved drugs is not their higher number of known targets *per se*, but an underlying factor correlated with it: the query drug and its top hits, which should include some of the chemical derivatives that eventually led to this drug, often have a lower overlap in terms of known targets. One contributing factor for a low overlap is that two similar chemical structures do not always have affinity for the same targets. There is abundant literature analysing these pathological cases known as activity cliffs (Medina-Franco, 2013). Furthermore, even the top hits might not be highly similar to the query molecule, although this issue will become less frequent as more molecules are included in chemogenomics databases. Another contributing factor is that some of the top hits could have been tested against a range of targets in other studies, which might not have included the drug and thus this molecule would not have been tested against the targets (a lower precision for this query molecule would be consequently obtained, as such targets would be perceived as false positives). Importantly, while these are not known targets of the drug, some are expected to become a known target once tested. In contrast, a molecule from the randomly-chosen set often has a larger overlap with its top hits (e.g. in set D, the predicted targets of 324 randomly-chosen molecules have precision=1, whereas those for just 35 randomly-chosen molecules have precision=0). The latter cases are likely to arise from a situation where a chemical series is investigated against a set of related targets to be later abandoned (Waring et al., 2015). This would explain the lower number of known targets and the smaller predictive errors for these sets.

### 3.4. How many known targets of the query molecule are typically missed?

Addressing this question is necessary to estimate how many discoveries are being missed by the ligand-centric method, but it has not been investigated with regards to employed query molecule. Figure 4 presents the results in terms of recall (middle plot). Looking at the recall boxplots, only about 10% of the targets are on average missed in the sets of random molecules (i.e. recall∼0.9), whereas the mean of missed targets for approved drugs is about 65%. A large part of these missed targets might be due to more intense research on the drug after approval than on its chemical derivatives, leading to many targets being tested in the former but not the latter.

On the other hand, the MCC boxplots (Figure 4 right plot) show the distribution of the total error across query molecules, with a high MCC necessarily meaning that the query molecule obtains low levels of both Type I and II errors. The latter occurs to most random molecules regardless of the activity threshold (almost 75% of these query molecules have MCCs higher than 0.6). In contrast, only a small proportion of approved drugs are in this category. Again, the performance hierarchy is D>B>C>A for both recall and MCC. Here, a higher number of known targets in the query molecules is also correlated with the difficulty of predicting their targets, but this is also explained by the different ways in which the query molecules and their hits were tested against targets.

### 3.5. Representative case studies

Section 3.2 analysed the NKT across query molecules. As discussed in section 3.3, the NKT of a molecule depends on its intrinsic polypharmacology, but also on how comprehensively the molecule has been tested across targets by the relevant scientific communities (we will call *observed polypharmacology* to the combination of these two factors). In the adopted TF method, the NPT of a molecule is given by the NKTs from its top 5 hits according to the dice score on MACCS fingerprints. Thus, the NPT for the query molecule depends in turn on the observed polypharmacology of each of these hits. In this section, we analyse two approved drugs representing cases where the difference in observed polypharmacology between the drug and its top hits are large in one direction (NKT>>NPT) or the other (NKT<<NPT).

The first case has nilotinib as the query molecule. Nilotinib was presented as a small-molecule selective tyrosine kinase inhibitor (Manley et al., 2010). However, we now know that this marketed drug has at least NKT=67 known molecular targets under 10μM, of which 14 are not kinases. In contrast, its top 5 hits collectively hit just nine targets (NPT=9). This is not surprising given the intense research interest in nilotinib as a targeted drug for the treatment of imatinib-resistant Chronic Myeloid Leukemia (Breccia and Alimena, 2010), but less so in its top hits from database molecules containing no drugs by construction. Figure 5A illustrates the proposed validation approach on nilotinib as the degree of overlap between the target spaces spanned by the query molecule and its top hits. Since predicted targets can only be either a true target or not, TP + FP = NPT. Likewise, known targets are either correctly predicted or not and thus TP + FN = NKT. As explained in section 2.4, the hit rate is given precision=TP/NPT, thus precision=0.89. In other words, the TF method retrospectively obtains an 89% hit rate for nilotinib (i.e. finding eight true targets of nilotinib in nine predicted targets). However, 59 known targets of nilotinib are missed by this method, as indicated by a low recall=TP/NKT of 0.12. This evidences that a method offering a high hit rate, while highly satisfying from a cost-effectiveness perspective, must be complemented by a high recall to be optimal.

**Figure 5.**
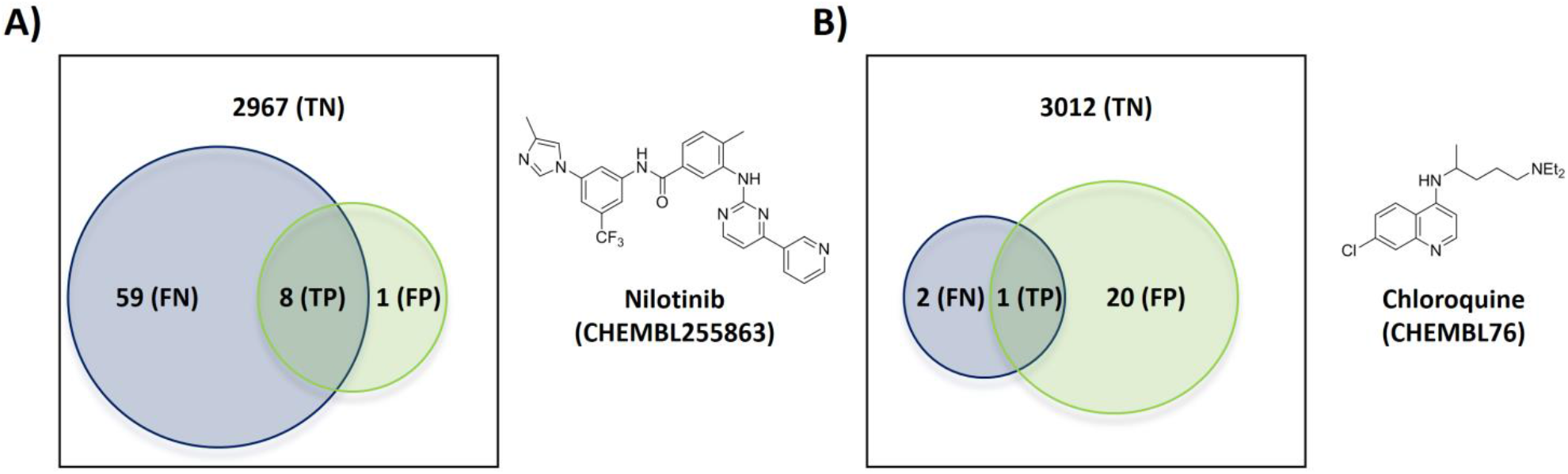
Overlap of the target spaces under 10μM spanned by the query molecule (blue circle representing its known targets) and its top hits (green circle representing predicted targets given by the known targets of its top 5 hits) according to the employed TF method. (**A**) Nilotinib as the query molecule. (**B**) Chloroquine as the query molecule.

Figure 5B shows the validation for the second case, which analyses the antimalarial agent chloroquine. NKT=3, NPT=21, precision=0.05 and recall=0.33 are obtained in these case. This represents a modest hit rate of just 5%, implying that a high experimental effort would have been associated to this discovery. However, the method obtains a higher recall with chloroquine than with nilotinib, which means that a lower proportion of known targets are being missed.

There are a total of 21 FP target predictions in both query molecules. However, none of these target-ligand pairs have actually been tested (i.e. no bioactivity associated to them in ChEMBL. Since they come from the most similar molecules to the query, it is likely that some of these predictions will result in the discovery of new targets of these query molecules once tested. For instance, the only FP of nilotinib is human GRM5 (metabotropic glutamate receptor 5; CHEMBL3227), which is a known target of the 4^th^ and 5^th^ most similar database molecules to nilotinib (CHEMBL2346729 and CHEMBL2346732, both with submicromolar affinity for this target). Another exciting prospect is one of the 20 unconfirmed FPs from chloroquine, human CCR4 (C-C chemokine receptor type 4; CHEMBL2414), which also a clinically-relevant target and also been predicted by two of the top 5 hits (CHEMBL194930 and CHEMBL195203, both with single-digit micromolar potency).

## 4. Conclusions

We have shown that ligand-centric techniques for TF are capable of considering up to thousands of targets more than target-centric techniques. This important advantage means that ligand-centric techniques have their niche in TF. We have also discussed the limitations of current benchmarks to test TF methods and consequently we have designed a new benchmark that overcomes them. Using the proposed benchmark, it has been possible to investigate how reliable are ligand-centric methods for TF depending on the employed query molecule. Despite the simplicity of the adopted method and owing to the wealth of data on which it relies, we have found that only five predicted targets will have to be tested in order to find two true targets with potency better than 1μM on average over marketed drugs. This level of performance is already useful for prospective applications and it is encouraging that there is plenty of scope for methodological improvement. The latter will be particularly needed to reduce the high number of false negatives, i.e. known targets that are currently missed by ligand-centric techniques. It is worth noting that, while this issue has not been investigated yet for target-centric techniques, the many targets not considered by this class of techniques are by construction false negatives of any molecule that hits them. We have argued that this drawback is hard to appreciate as target-centric techniques only report predictive performance achieved on the typically much smaller set of considered targets.

The results for the set of randomly-selected molecules used as a control group are substantially better than those for approved drugs. We have discussed how the different way in which targets are tested against the query molecules and their top hits is the primary reason for this marked difference. Since approved drugs have been presumably tested against many more targets, we consider that their performance level is more realistic than that of the control set. Interestingly, we have identified an average of eight known targets under 10μM in approved drugs, which reinforces the notion that polypharmacology is a common and strong event. Lastly, high performance variability across query molecules has been observed in all cases. Thus, a promising avenue for future research consists in investigating which features make the target-ligand pair more difficult to predict in order to assign a confidence score to each prediction.

## Author contributions

P.J.B. conceived the study and wrote the manuscript. A.P. implemented the software and carried out the numerical experiments with the assistance of C.C.D. All authors reviewed the manuscript.

## Acknowledgments

This work has been carried out thanks to the support of the A^*^MIDEX grant (n° ANR-11-IDEX-0001-02) funded by the French Government «Investissements d’Avemr» program awarded to P.J.B.

## Additional information

The authors declare no competing financial interests.

## References

AbdulHameed, M. D. M., Chaudhury, S., Singh, N., Sun, H., Wallqvist, A., and Tawa, G. J.(2012). Exploring polypharmacology using a ROCS-based target fishing approach. J. Chem. Inf. Model. 52, 492–505. doi:10.1021/ci2003544.

Ain, Q. U., Aleksandrova, A., Roessler, F. D., and Ballester, P. J. (2015). Machine-learning scoring functions to improve structure-based binding affinity prediction and virtual screening. Wiley Interdiscip. Rev. Comput. Mol. Sci. 5, 405–424. doi:10.1002/wcms.1225.

Armstrong, M., Morris, G., Finn, P., Sharma, R., Moretti, L., Cooper, R., et al. (2010). ElectroShape: fast molecular similarity calculations incorporating shape, chirality and electrostatics. J. Comput. Aided. Mol. Des. doi:10.1007/s10822-010-9374-0.

Ballester, P. J. (2011). Ultrafast shape recognition: method and applications. Future Med. Chem. 3, 65–78.

Ballester, P. J., and Richards, W. G. (2007). Ultrafast shape recognition for similarity search in molecular databases. Proc. R. Soc. A Math. Phys. Eng. Sci. 463, 1307–1321. doi:10.1098/rspa.2007.1823.

Bento, A. P., Gaulton, A., Hersey, A., Bellis, L. J., Chambers, J., Davies, M., et al. (2014). The ChEMBL bioactivity database: an update. Nucleic Acids Res. 42, D1083–90. doi:10.1093/nar/gkt1031.

Breccia, M., and Alimena, G. (2010). Nilotinib: a second-generation tyrosine kinase inhibitor for chronic myeloid leukemia. Leuk. Res. 34, 129–34. doi:10.1016/j.leukres.2009.08.031.

Cereto-Massagué, A., Ojeda, M. J., Valls, C., Mulero, M., Pujadas, G., and Garcia-Vallve, S. (2015). Tools for in silico target fishing. Methods 71, 98–103. doi:10.1016/j.ymeth.2014.09.006.

ChEMBL_20 release Available at: ftp://ftp.ebi.ac.uk/pub/databases/chembl/ChEMBLdb/releases/chembl_20/. [Accessed March 4, 2015].

Cheng, T., Pan, Y., Hao, M., Wang, Y., and Bryant, S. H. (2014). PubChem applications in drug discovery: a bibliometric analysis. Drug Discov. Today 19, 1751–6. doi:10.1016/j.drudis.2014.08.008.

Cortés-Cabrera, A., Morris, G. M., Finn, P. W., Morreale, A., and Gago, F. (2013). Comparison of ultra-fast 2D and 3D ligand and target descriptors for side effect prediction and network analysis in polypharmacology. Br. J. Pharmacol. 170, 557–67. doi:10.1111/bph.12294.

Durant, J. L., Leland, B. A., Henry, D. R., and Nourse, J. G. Reoptimization of MDL keys for use in drug discovery. J. Chem. Inf. Comput. Sci. 42, 1273–80. Available at: http://www.ncbi.nlm.nih.gov/pubmed/12444722 [Accessed June 19, 2015].

Füllbeck, M., Dunkel, M., Hossbach, J., Daniel, P. T., and Preissner, R. (2009). Cellular Fingerprints: A Novel Approach Using Large-Scale Cancer Cell Line Data for the Identification of Potential Anticancer Agents. Chem. Biol. Drug Des. 74, 439–448. doi:10.1111/j.1747-0285.2009.00883.x.

Gao, Z., Li, H., Zhang, H., Liu, X., Kang, L., Luo, X., et al. (2008). PDTD: a web-accessible protein database for drug target identification. BMC Bioinformatics 9, 104. doi:10.1186/1471-2105-9-104.

Gfeller, D., Grosdidier, A., Wirth, M., Daina, A., Michielin, O., and Zoete, V. (2014). SwissTargetPrediction: a web server for target prediction of bioactive small molecules. Nucleic Acids Res., gku293–. doi:10.1093/nar/gku293.

Holbeck, S. L., Collins, J. M., and Doroshow, J. H. (2010). Analysis of FDA-Approved AntiCancer Agents in the NCI60 Panel of Human Tumor Cell Lines. Mol. Cancer Ther. 9, 1451–1460. doi:10.1158/1535-7163.MCT-10-0106.

Huang, H., Zhang, P., Qu, X. A., Sanseau, P., and Yang, L. (2014). Systematic prediction of drug combinations based on clinical side-effects. Sci. Rep. 4, 7160. doi:10.1038/srep07160.

Keiser, M. J., Roth, B. L., Armbruster, B. N., Ernsberger, P., Irwin, J. J., and Shoichet, B. K. (2007). Relating protein pharmacology by ligand chemistry. Nat. Biotechnol. 25, 197–206. doi:10.1038/nbt1284.

Keiser, M. J., Setola, V., Irwin, J. J., Laggner, C., Abbas, A. I., Hufeisen, S. J., et al. (2009). Predicting new molecular targets for known drugs. Nature 462, 175–181. doi:10.1038/nature08506.

Koutsoukas, A., Lowe, R., Kalantarmotamedi, Y., Mussa, H. Y., Klaffke, W., Mitchell, J. B. O., et al. (2013). In silico target predictions: defining a benchmarking data set and comparison of performance of the multiclass Naïve Bayes and Parzen-Rosenblatt window. J. Chem. Inf. Model. 53, 1957–66. doi:10.1021/ci300435j.

van Laarhoven, T., Nabuurs, S. B., and Marchiori, E. (2011). Gaussian interaction profile kernels for predicting drug-target interaction. Bioinformatics 27, 3036–43. doi:10.1093/bioinformatics/btr500.

Lamdrum, G. RDKit: Open-source cheminformatics. Available at: http://www.rdkit.org/ [Accessed April 3, 2015].

Lavecchia, A., and Cerchia, C. (2015). In silico methods to address polypharmacology: Current status, applications and future perspectives. Drug Discov. Today 21, 288–298. doi:10.1016/j.drudis.2015.12.007.

Lee, J., and Bogyo, M. (2013). Target deconvolution techniques in modern phenotypic profiling. Curr. Opin. Chem. Biol. 17, 118–126. doi:10.1016/j.cbpa.2012.12.022.

Liu, X., Xu, Y., Li, S., Wang, Y., Peng, J., Luo, C., et al. (2014). In Silico target fishing: addressing a “Big Data” problem by ligand-based similarity rankings with data fusion. J. Cheminform. 6, 33. doi:10.1186/1758-2946-6-33.

Manley, P. W., Stiefl, N., Cowan-Jacob, S. W., Kaufman, S., Mestan, J., Wartmann, M., et al. (2010). Structural resemblances and comparisons of the relative pharmacological properties of imatinib and nilotinib. Bioorg. Med. Chem. 18, 6977–86. doi:10.1016/j.bmc.2010.08.026.

Martínez-Jiménez, F., Papadatos, G., Yang, L., Wallace, I. M., Kumar, V., Pieper, U., et al. (2013). Target prediction for an open access set of compounds active against Mycobacterium tuberculosis. PLoS Comput. Biol. 9, e1003253. doi:10.1371/journal.pcbi.1003253.

Medina-Franco, J. L. (2013). Activity Cliffs: Facts or Artifacts? Chem. Biol. Drug Des. 81, 553–556. doi:10.1111/cbdd.12115.

Mestres, J., Gregori-Puigjané, E., Valverde, S., and Solé, R. V (2009). The topology of drug-target interaction networks: implicit dependence on drug properties and target families. Mol. Biosyst. 5, 1051–1057. doi:10.1039/b905821b.

Mugumbate, G., Abrahams, K. A., Cox, J. A. G., Papadatos, G., van Westen, G., Lelièvre, J., et al. (2015). Mycobacterial dihydrofolate reductase inhibitors identified using chemogenomic methods and in vitro validation. PLoS One 10, e0121492. doi:10.1371/journal.pone.0121492.

Nettles, J. H., Jenkins, J. L., Bender, A., Deng, Z., Davies, J. W., and Glick, M. (2006). Bridging chemical and biological space: “target fishing” using 2D and 3D molecular descriptors. J. Med. Chem. 49, 6802–10. doi:10.1021/jm060902w.

Nigsch, F., Bender, A., Jenkins, J. L., and Mitchell, J. B. O. (2008). Ligand-target prediction using Winnow and naive Bayesian algorithms and the implications of overall performance statistics. J. Chem. Inf. Model. 48, 2313–25. doi:10.1021/ci800079x.

Papadatos, G., Davies, M., Dedman, N., Chambers, J., Gaulton, A., Siddle, J., et al. (2015). SureChEMBL: a large-scale, chemically annotated patent document database. Nucleic Acids Res., gkv1253–. doi:10.1093/nar/gkv1253.

Rogers, D., and Hahn, M. (2010). Extended-connectivity fingerprints. J. Chem. Inf. Model. 50, 742–54. doi:10.1021/ci100050t.

Schneider, G. (2010). Virtual screening: an endless staircase? Nat Rev Drug Discov 9, 273–276. doi:10.1038/nrd3139.

Schomburg, K. T., and Rarey, M. (2014). Benchmark data sets for structure-based computational target prediction. J. Chem. Inf. Model. 54, 2261–74. doi:10.1021/ci500131x.

Speck-Planche, A., and Cordeiro, M. N. D. S. (2015). Multi-Target QSAR Approaches for Modeling Protein Inhibitors. Simultaneous Prediction of Activities Against Biomacromolecules Present in Gram-Negative Bacteria. Curr. Top. Med. Chem. 15, 1801–13. Available at: http://www.ncbi.nlm.nih.gov/pubmed/25961517 [Accessed February 18, 2016].

Sukumar, N., and Das, S. (2011). Current trends in virtual high throughput screening using ligand-based and structure-based methods. Comb. Chem. High Throughput Screen. 14, 872–888. Available at: http://www.ncbi.nlm.nih.gov/pubmed/21843144.

Ursu, A., and Waldmann, H. (2015). Hide and seek: Identification and confirmation of small molecule protein targets. Bioorg. Med. Chem. Lett. 25, 3079–86. doi:10.1016/j.bmcl.2015.06.023.

Wang, L., Ma, C., Wipf, P., Liu, H., Su, W., and Xie, X.-Q. (2013). TargetHunter: an in silico target identification tool for predicting therapeutic potential of small organic molecules based on chemogenomic database. AAPS J. 15, 395–406. doi:10.1208/s12248-012-9449-z.

Waring, M. J., Arrowsmith, J., Leach, A. R., Leeson, P. D., Mandrell, S., Owen, R. M., et al. (2015). An analysis of the attrition of drug candidates from four major pharmaceutical companies. Nat. Rev. Drug Discov. 14, 475–486. doi:10.1038/nrd4609.

Willett, P. (2014). The Calculation of Molecular Structural Similarity: Principles and Practice. Mol. Inform. 33, 403–413. doi:10.1002/minf.201400024.

Yu, H., Chen, J., Xu, X., Li, Y., Zhao, H., Fang, Y., et al. (2012). A systematic prediction of multiple drug-target interactions from chemical, genomic, and pharmacological data. PLoS One 7, e37608. doi:10.1371/journal.pone.0037608.

Zanni, R., Gálvez-Llompart, M., Gálvez, J., and García-Domenech, R. (2014). QSAR multi-target in drug discovery: a review. Curr. Comput. Aided. Drug Des. 10, 129–36. Available at: http://www.ncbi.nlm.nih.gov/pubmed/24724898 [Accessed February 18, 2016].

